# Desert hedgehog enhances endothelial resilience and prevents atherosclerosis by mitigating PAI-1 signaling

**DOI:** 10.64898/2026.06.02.729715

**Authors:** Iqra Ilyas, Zhihua Wang, Meiming Su, Hui Jiang, Zhidan Zhang, Fang Zhu, Yun Fang, Li Wang, Yu Huang, Marie Ange Renault, Jaroslav Pelisek, Giovanni G. Camici, Hanjoong Jo, Paul C. Evans, Stefan Offermanns, Clint L. Miller, Floor B H van der Zalm, Sander W. van der Laan, Lars Maegdefesse, Yulei Zhang, Qingqing Wu, Jizhou Zhang, Bin Zhou, Tengchuan Jin, Suowen Xu, Jianping Weng

## Abstract

**BACKGROUND:** Atherosclerotic cardiovascular disease remains the leading cause of death worldwide. Most current pharmacotherapies target conventional risk factors that promote atherosclerosis (e.g., hyperlipidemia) rather than intrinsic resilience factors that protect against atherosclerosis in the face of risk factors. Here, we investigated the role of desert hedgehog (*DHH*), a canonical ligand of the hedgehog signaling pathway, as a novel resilience factor that restrains endothelial mesenchymal transition (EndoMT) and protects against atherosclerosis.

**METHODS:** Single-cell RNA sequencing (scRNA-seq) was performed on atheroprone and atheroprotective regions of the *ApoE* knockout mouse aorta to identify mechanoresponsive genes associated with atherosclerosis. Endothelial cell-specific *Dhh* knockout mice were subjected to partial carotid ligation and hypercholesterolemic conditions to investigate the role of endothelial *Dhh* in atherosclerosis progression. scRNA-seq, bulk RNA sequencing, endothelial lineage tracing, immunoprecipitation-coupled mass spectrometry, and surface plasmon resonance were used to investigate the role and mechanism of *DHH* in EndoMT. Pharmacological interventions and recombinant DHH administration were performed *in vivo* to evaluate therapeutic potential of DHH targeting. DHH expression was also examined in human atherosclerotic arteries and serum samples from patients with coronary artery disease.

**RESULTS:** DHH protein expression was enriched in arterial endothelium from mice, porcine, and humans. However, DHH expression was significantly reduced in atherosclerotic arteries and serum from patients with coronary artery disease. scRNA-seq of atheroprone and atheroresistant region of mouse aorta identified *Dhh* as a novel mechanoresponsive gene enriched in aortic regions exposed to unidirectional laminar flow. Endothelial cell-specific *Dhh* knockout (*Dhh*^ecKO^) mice exhibited increased atherosclerotic lesion area, large necrotic cores, and reduced collagen content following partial carotid ligation. Similarly, under hypercholesterolemic conditions, both male and female *Dhh*^ecKO^ mice showed aggravated atherosclerosis progression. scRNA-seq of *Dhh*^ecKO^ mouse aortas revealed an increased proportion of endothelial cells undergoing mesenchymal transition, indicating enhanced EndoMT. These findings were corroborated by bulk RNA-sequencing of *DHH* depleted HUVECs and endothelial lineage tracing in inducible *Dhh*^ecKO^ mice. Mechanistically, DHH directly interacted with plasminogen activator inhibitor type 1 (PAI-1) and suppressed PAI-1-induced EndoMT. PAI-1 promoted EndoMT in ECs through activation of canonical TGF-β signaling (SMAD2/3) and noncanonical AKT/ERK1/2 signaling via interaction with low-density lipoprotein receptor-related protein (LRP1). Neutralization of PAI-1 or inhibition of LRP1, AKT/ERK1/2, or SMAD3 signaling abolished *DHH* deficiency-induced EndoMT. DHH competitively inhibited PAI-1 binding to LRP1, thereby attenuating downstream pro-EndoMT signaling. Intriguingly, treatment with PAI-1 inhibitor TM5275 mitigated endothelial *Dhh* deficiency induced EndoMT *in vivo*. Of translational relevance, recombinant mouse DHH protein administration reduced atherosclerosis progression, stabilized plaque, and decreased the expression of EndoMT markers in *ApoE* knockout mice.

**CONCLUSIONS:** Desert hedgehog (DHH) is an intrinsic endothelial cell-enriched resilience factor that protects against EndoMT and atherosclerosis by preventing PAI-1 signaling. The present study implicates endothelial DHH as a potential therapeutic target for atherosclerotic cardiovascular disease.

**Graphical abstract:** 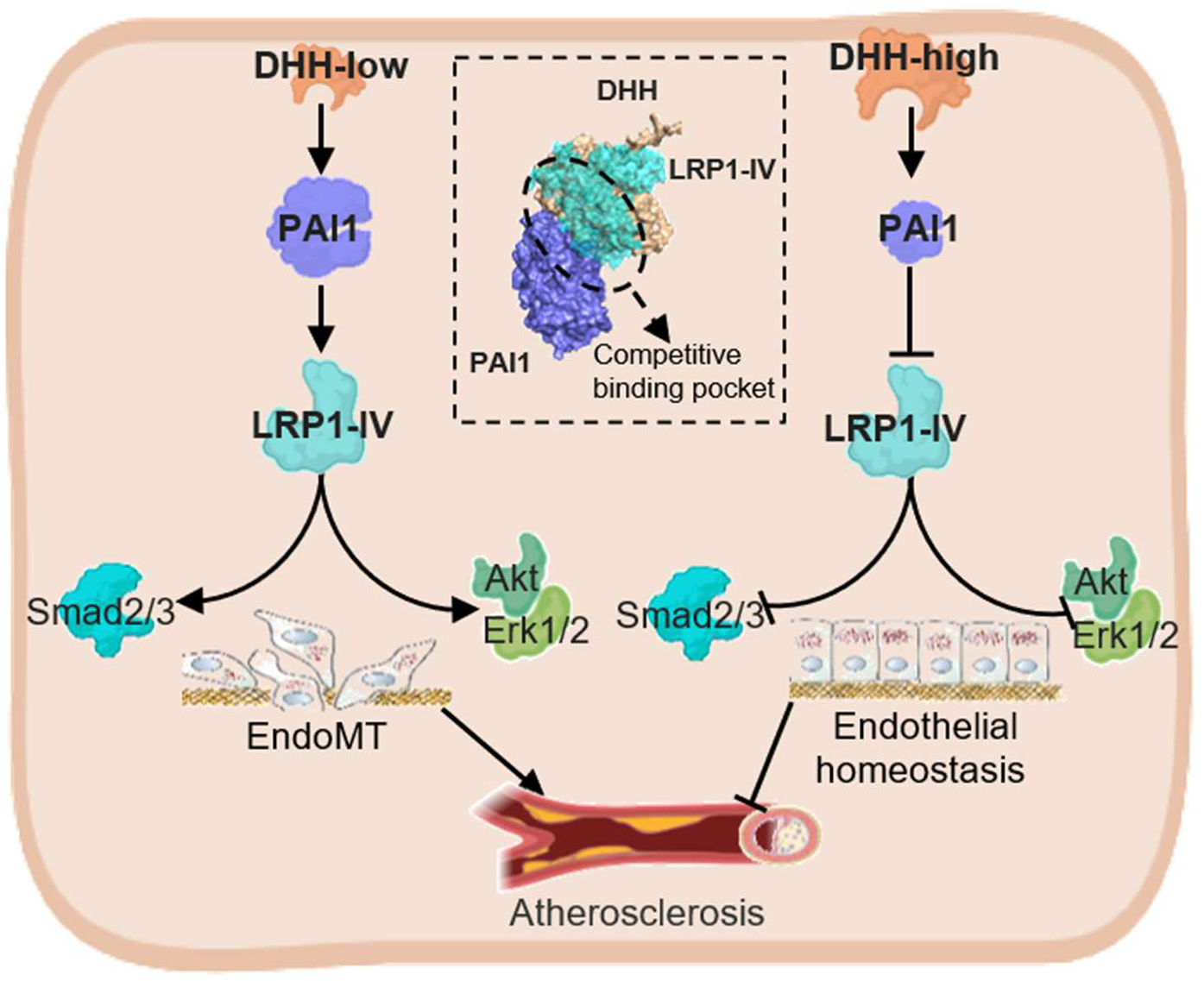

Desert hedgehog (DHH) is an intrinsic endothelial cell-enriched resilience factor that protects against EndoMT and atherosclerosis by preventing PAI-1 binding to LRP1 and downstream AKT/ERK, as well as SMAD2/3 signaling.

**Clinical Perspective:** *What Is New?:*

- *DHH* was identified as a flow-responsive resilience gene that is downregulated by disturbed flow in endothelial cells.
- Endothelial-specific deletion of *Dhh* promoted endothelial-to-mesenchymal transition (EndoMT) and atherosclerosis.
- DHH directly interacts with PAI-1 and preclude PAI-1 mediated pro-EndoMT signaling.

*What Are the Clinical Implications?:*

- DHH maintains endothelial homeostasis during atherosclerosis.
- Targeting endothelial DHH-PAI-1 axis may represent a potential therapeutic strategy to reduce EndoMT and limit plaque progression.
- Lower circulating DHH level may serve as a potential biomarker of endothelial dysfunction and plaque vulnerability in atherosclerotic disease.

## INTRODUCTION

Atherosclerosis is a progressive vascular disease characterized by the accumulation of fatty and fibrous material within the arterial wall, leading to plaque formation.^1, 2^ It is a leading cause of coronary artery disease (CAD), myocardial infarction, and ischemic stroke, contributing to significant morbidity and mortality worldwide.^3, 4^ Traditionally, intensive research efforts have focused on developing pharmacotherapies that target disease-causing risk factors (e.g. hyperlipidemia). Nowadays, identifying the molecular underpinnings of endothelial resilience in the face of existing risk factors yields a paradigm shift toward reducing residual cardiovascular risk.^5–7^ Disturbed hemodynamic forces are biomechanical risk factors for atherosclerosis development.^8–10^ Atherosclerotic plaques preferentially develop in arterial regions exposed to disturbed flow (DF), such as vascular bifurcations and vascular bends.^11–13^ Conversely, arterial regions exposed to unidirectional laminar flow (UF), such as straight and non-branching segments, are relatively protected from lesion formation.^8, 10, 12, 13^ In DF regions, endothelial cells (ECs) experience oscillatory shear stress, which triggers endothelial dysfunction, including inflammation, cell proliferation, apoptosis, increased permeability, and endothelial-to-mesenchymal transition (EndoMT).^13–16^ In contrast, UF promotes ECs alignment, homeostasis, and atheroprotective transcriptional reprogramming.^12, 13, 17^ Therefore, UF-mediated mechanotransduction is a major contributor to endothelial resilience against atherogenic stress.

EndoMT plays a pivotal role in multiple aspects of atherosclerotic disease progression.^18^ It is an intricate process of endothelial reprogramming and redifferentiation characterized by endothelial detachment, migration, partial or complete loss of endothelial characteristics, and acquisition of mesenchymal features.^19, 20^ EndoMT is primarily defined by morphological changes and altered expression of lineage-specific markers. For example, endothelial markers such as CD31, VE-cadherin and ICAM2 decrease, whereas mesenchymal markers such as FN1 and α-SMA increase during EndoMT.^21–23^ EndoMT has also been associated with increased expression of plasminogen activator inhibitor 1 (PAI-1), which can itself further promote EndoMT,^24^ thereby creating a vicious cycle. EndoMT is closely associated with increased plaque burden^25^ and plaque instability in both mice and humans.^20, 26^ Fate-mapping studies suggests that >30% of luminal ECs in mice with advanced lesions express EndoMT-associated markers.^20^ Similarly, approximately 70% of ECs in severe human coronary plaques exhibit EndoMT-related marker expression.^20^ Numerous stimuli, including growth factors, cytokines, inflammation, oxidative stress, and DF can induce EndoMT, and several signaling pathways, particularly transforming growth factor-β (TGF-β) signaling, have been implicated in this process during atherosclerosis.^20, 25, 27^ Further understanding the mechanisms governing EndoMT is therefore crucial for developing therapeutic strategies to prevent or reverse atherosclerosis.

DHH is a canonical ligand of the hedgehog signaling pathway which has been implicated in diverse developmental and cellular processes.^28, 29^ Emerging evidence indicates that hedgehog signaling regulates vascular development, angiogenesis, and vascular permeability.^30–37^ DHH down-regulation has been associated with cardiovascular risk factors, including obesity, diabetes mellitus, and aging in both rodents and humans.^38^ However, the precise role of DHH in regulating EndoMT and atherosclerosis remains unknown.

Here, by leveraging single-cell RNA sequencing (scRNA-seq) of distinct arterial regions from atheroprone apolipoprotein E knockout (*ApoE* KO) mouse aortas, we identified *Dhh* as a novel mechanoresponsive gene highly expressed in regions exposed to UF. We further found that endothelial-specific deletion of *Dhh* promoted EndoMT and accelerated atherosclerosis in mice. Conversely, administration of recombinant mouse DHH (rDHH) protein to *ApoE* KO mice mitigated EndoMT and reduced lesion formation. Consistent with potential clinical relevance, DHH protein levels were significantly reduced in both atherosclerotic arteries and serum from patients with coronary artery disease. Collectively, these findings identify *DHH* as an intrinsic endothelial resilience factor that restrains atherosclerosis by limiting EndoMT and suggests that *DHH* may represent a novel therapeutic target in vascular diseases driven by EndoMT.

## METHODS

### Data Availability

A detailed description of the methods and supporting data is available in the <u>Supplemental Mate-rial</u>. Related data supporting this study’s findings can be obtained from the corresponding author upon reasonable request. The raw scRNA-seq and RNA-seq datasets have been submitted to the China National Center for Bioinformation / Beijing Institute of Genomics, Chinese Academy of Sciences, under accession numbers CRA016472 and CRA016471, respectively. These datasets are publicly accessible at https://ngdc.cncb.ac.cn/gsa. Additionally, the mass spectrometry proteomics data have been deposited in the ProteomeXchange Consortium via the iProX partner repository, with the dataset identifier PXD052429, available at https://proteomecentral.proteomexchange.org.

### Human samples

Human serum and tissue samples were collected from The First Affiliated Hospital of the University of Science and Technology of China (USTC). The study received approval from the Institutional ethics committee (Approval No: 2021KY-089). Atherosclerotic plaque tissue samples were obtained from the coronary arteries of heart transplant recipients with heart failure following myocardial infarction (n = 3), while healthy tissue samples were obtained from the coronary arteries of heart transplant donors (n = 3). Serum samples from healthy individuals were collected from a health examination center (n = 10), while serum from coronary artery disease (CAD) patients was obtained from the cardiac surgery department (n = 10). Serum was used for ELISA assays, and coronary artery tissues were employed for immunofluorescence staining. This study was performed in accordance with the Helsinki Declaration guidelines, with written informed consent obtained from all participants. Demographic characteristics of the human samples used for immuno-fluorescence staining (n=3) and ELISA (n=10) can be found in our previously published study.^39^

### Experimental Animals

All animal experimental procedures used in this study were conducted in accordance with the NIH “Guide for the Care and Use of Laboratory Animals”. Approval for the animal experiments was granted by the Laboratory Animal Ethics Committee of the USTC (Approval number for mouse experiments is USTCACUC27120124059 and approval number for pig experiments is 2023-N(A)-0172). Endothelial cell-specific *Dhh* knockout mice, endothelial lineage tracing mice, and *ApoE* KO mice were used to explore the role of *Dhh* in atherosclerosis progression and EndoMT. Procedures for animal experiments conform to the ARRIVE (Animal Research: Reporting of In Vivo Experiments) guidelines 2.0.

### Statistical Analysis

Statistical analyses were carried out using GraphPad Prism version 8.0.1. Data are presented as mean ± SEM unless otherwise stated. The Shapiro-Wilk test was used to assess data normality. For comparisons between two groups, an unpaired two-tailed Student’s t-test was applied for normally distributed data with equal variances, with Welch’s correction used for normally distributed data when variances were unequal. For nonnormal distribution data, a two-tailed nonparametric Mann-Whitney *U* test was used. When analyzing 3 or more groups, 1-way ANOVA followed by Bonferroni post hoc test was used. When analyzing more than 2 groups with 2 or more variables, a 2-way ANOVA followed by Bonferroni post hoc test was used. Statistical significance was defined as P < 0.05; all *p*-values are reported in the figures. Each experiment was conducted at least 3 times to ensure reliable and reproducible results, with the sample size (n) specified in the corresponding figure legends.

## RESULTS

### scRNA-seq analysis of *ApoE* KO mouse aorta identifies *Dhh* as a novel athero-relevant and mechanoresponsive gene

To discover new athero-relevant and mechanosensitive genes that may contribute to atherosclerosis progression, we performed scRNA-seq analysis of two anatomically distinct regions of the aorta, namely the aortic arch (AA) and descending thoracic aorta (dTA), from *ApoE* KO mice following 12 weeks of high-cholesterol diet (HCD) feeding (Fig. 1A). Atheroprone AA and athero-protective TA regions of the aorta experience DF and UF patterns, respectively, making them ideal contrasting models for studying flow-dependent factors *in vivo*. Following preprocessing and quality-control filtering, 12,698 cells were retained for downstream analysis (Fig. S1A and S1B). Using the Seurat platform for single-cell transcriptomic analysis, we initially identified 18 distinct cell clusters (Fig. S1C). Based on established marker genes for major vascular and immune cell populations within the mouse aorta (Fig. 1B), these clusters were subsequently annotated into 7 cell types, including vascular smooth muscle cells (VSMCs), fibroblasts, macrophages, ECs, T lymphocytes (T cells), natural killer cells (NK cells), and B lymphocytes (B cells) (Fig. 1C and Fig. S1D). Further analysis of EC-enriched genes demonstrated marked differences in the expression of endothelial-associated markers, including *Klf2*, *Tek*, *Heg1*, and *Serpine1,* between the AA and TA regions (Fig. 1D). These findings are consistent with regional endothelial heterogeneity associated with distinct hemodynamic environments in the mouse aorta.

**Fig. 1.**
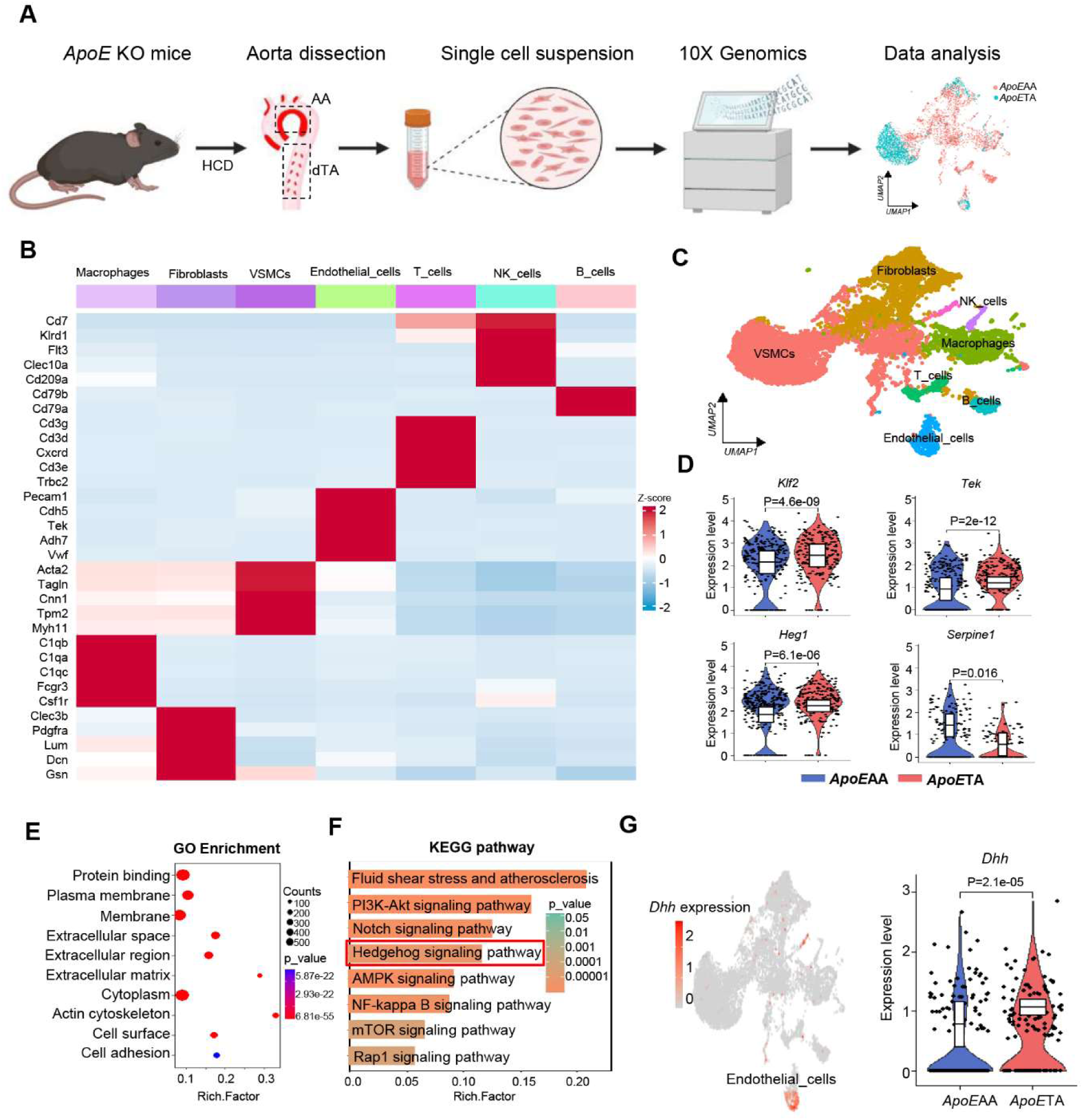
Identification of *Dhh* as a novel athero-relevant and mechanoresponsive gene via scRNA seq analysis. **(A**) Workflow diagram of scRNA-seq analysis of aortic arch (AA) and descending thoracic aorta (dTA) of *ApoE* KO mice fed a high cholesterol diet for 12-weeks. (**B**) Heatmap displays the top 5 markers for each cluster in scRNA-seq. A total of 22,847 cells were divided into 19 clusters, which were ultimately identified as 7 distinct cell types. (**C**) UMAP plot of scRNA for all samples, colored according to cell type. (**D**) Violin plot compares the relative expression levels of representative genes in ECs between *ApoE*AA and *ApoE*TA. (**E**) Bubble chart displays the GO enrichment analysis results of differentially expressed genes between *Apo-E*TA and *ApoE*AA. (**F**) Bar chart displays the KEGG signaling pathway analysis results of differentially expressed genes between *ApoE*TA and *ApoE*AA. (**G**) UMAP plot and violin plot demonstrate the specific expression of the *Dhh* gene in ECs, as well as significant expression differences between *ApoE*TA and *ApoE*AA. Statistical analysis was performed using wilcox.test for **D** and **G**.

To further characterize endothelial transcriptional changes between the *ApoE*AA and *Apo-E*TA groups, we performed Gene Ontology (GO) enrichment analysis of differentially expressed genes in ECs. GO analysis identified enrichment of biological processes related to protein binding, cell membrane organization, and extracellular matrix adhesion (Fig. 1E). In parallel, Kyoto Encyclopedia of Genes and Genomes (KEGG) pathway analysis revealed enrichment of pathways associated with fluid shear stress and atherosclerosis, PI3K-AKT signaling, Notch signaling, and hedgehog signaling (Fig. 1F). Genes contributed to individual KEGG pathway enrichments are listed in Supplementary Table S1. Five genes contributed to the enrichment of the hedgehog signaling pathway, namely *Dhh, Ptch1, Ccnd1, Ccnd2,* and *Cul1*. To further evaluate endothelial hedgehog signaling profiles, we examined the expression of key hedgehog pathway genes in ECs from the AA and TA regions, and the relative expression patterns are presented as a heatmap in Figure S2A. Interestingly, *Dhh* gene showed major differential expression between endothelial populations from the AA and TA regions (Fig. S2A and 1G), suggesting its potential association with region-specific endothelial phenotypes in the mouse aorta.

### Disturbed flow downregulates *DHH* expression in ECs

To investigate the potential role of *DHH* in vascular biology and atherosclerosis, we first examined the expression of hedgehog ligands across multiple vascular cell types, including human umbilical vein endothelial cells (HUVECs), human coronary artery endothelial cells (HCAECs), human pulmonary microvascular endothelial cells (HPMECs), human aortic smooth muscle cells (HASMC) and THP1-derived macrophages. Among the three hedgehog ligands (namely *DHH,* Indian hedgehog (*IHH*) and sonic hedgehog (*SHH*)), *DHH* showed significantly higher mRNA expression in human ECs from three different vascular beds (Fig. S3A). In contrast, *DHH* expression was comparatively lower in other atherorelevant cell types, including HASMCs and THP1-derived macrophages (Fig. S3B), suggesting that *DHH* is the primary hedgehog ligand expressed by ECs. To further assess endothelial *DHH* expression in human atherosclerotic disease, we analyzed a publicly available scRNA-seq dataset of calcified plaques (GSE159677), and observed enriched *DHH* expression in plaque-associated EC populations (Fig. S3C). Consistent with these transcriptomic findings, immunofluorescence (IF) staining of coronary arteries from mice, pigs, and humans demonstrated prominent DHH expression within the endothelial layer, where it colocalized with endothelial markers including CD31 and vWF (Fig. S3D-S3F).

Previous studies have identified *DHH* as a downstream target of KLF2.^37^ To examine whether *DHH* expression is associated with KLF2 activation in ECs, we treated HUVECs with atorvastatin, which is known to enhance KLF2 expression and activity. Atorvastatin increased *DHH* mRNA expression in HUVECs (Fig. S4), supporting an association between KLF2 activation and *DHH* expression in ECs.

Because our scRNA-seq analysis suggested flow-dependent regulation of *DHH*, we next evaluated *DHH* expression under distinct hemodynamic conditions. Analysis of two publicly available datasets (GSE20739, GSE160611) demonstrated that *DHH* mRNA expression was lower in ECs exposed to DF compared with UF (Fig. 2A and 2B). To determine whether *Dhh* expression is similarly regulated by blood flow patterns *in vivo*, we analyzed *Dhh* expression in distinct regions of the mouse aorta exposed to different hemodynamic environments, namely the inner curvature of the aortic arch (AA; DF region) and the descending thoracic aorta (TA; UF region) (Fig. 2C). *Dhh* mRNA expression was significantly lower in the AA compared with the TA (Fig. 2C). Consistent with these findings, *en-face* IF staining of mouse aortic endothelium demonstrated reduced DHH protein expression in the AA relative to the TA (Fig. 2D).

**Fig. 2.**
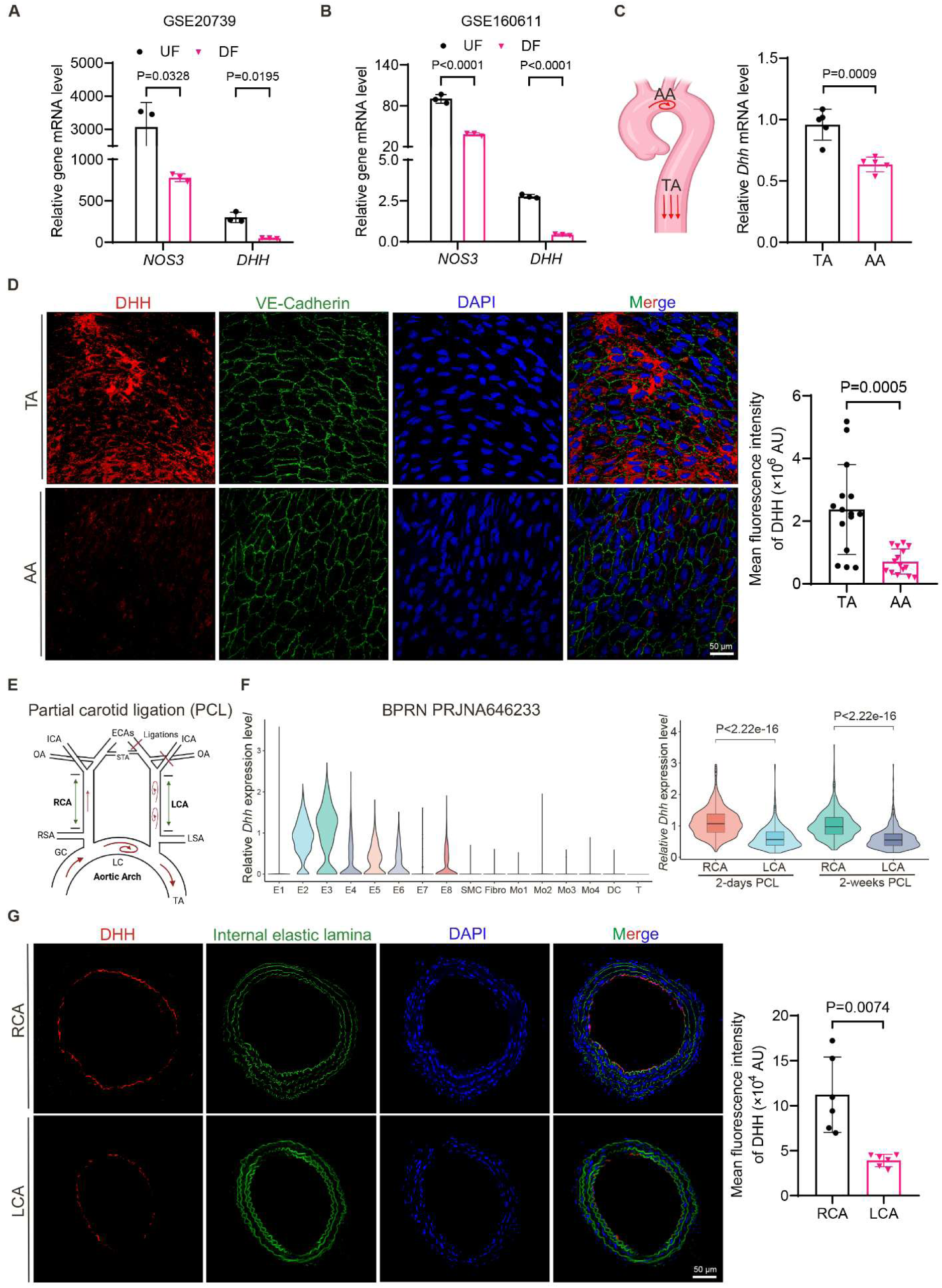
*Dhh* expression is downregulated by disturbed flow *in vitro* and *in vivo.* **(A**) The ex-pression of *NOS3* and *DHH* in HUVECs subjected to UF and DF in a public microarray dataset (GSE20739; n=3). (**B**) The expression of *NOS3* and *DHH* in HAECs subjected to UF and DF in a public RNA sequencing dataset (GSE160611; n=3). (**C**) RT-qPCR analysis of the mRNA levels of *Dhh* in the descending TA and AA regions of the C57BL/6J mouse aorta (n=5). (**D**) Representative *en-face* immunostaining of DHH (red), VE-Cadherin (green) and DAPI (blue) in the descend-ing TA and AA regions of the C57BL/6J mouse aorta (n=5, at least 3 different areas were analyzed per mouse). (**E)** Schematic diagram of partial carotid ligation (PCL) surgery in the mouse model. (**F**) The expression of *DHH* in EC-enriched bulk RNAs from the right carotid artery (RCA) and left carotid artery (LCA) after 2-days and 2-weeks of PCL in a public scRNA-seq dataset. (NCBI BioProject repository accession number is PRJNA646233; n=10). (**G**) Representative immunostaining images of DHH (red), auto-fluorescence of the internal elastic lamina (green) and DAPI (blue) in the RCA and LCA of male C57BL/6J mice after 3-weeks of PCL (n=6). All data are presented as mean ± SEM. Statistical analysis was performed using unpaired two-tailed Stu-dent t-test for **A**, **B**, **C**, **D** and **G**, and wilcox.test for right panel of **F**.

To further investigate the *Dhh* response to acute blood flow changes, we analyzed a publicly available dataset^40^ from a mouse partial carotid ligation (PCL) model, in which DF is induced in the left carotid artery (LCA), whereas the right carotid artery (RCA) remains exposed to UF (scheme shown in Fig. 2E). At baseline, *Dhh* expression was higher in ECs compared to other vascular cell populations (Fig. 2F). Following PCL, *Dhh* expression was significantly reduced in the LCA compared with the RCA at 48 hours and was further decreased after 2 weeks (Fig. 2F, right panel). Similar to these transcriptomic findings, IF staining demonstrated lower DHH protein expression in the LCA compared with the RCA following PCL (Fig. 2G). Collectively, these findings suggest that *DHH* is a flow-responsive gene in endothelial cells that is downregulated under DF conditions associated with atheroprone vascular regions.

### EC-specific deletion of *Dhh* aggravates atherosclerosis development in mice

To investigate the EC-specific role of *Dhh* in atherosclerosis *in vivo*, we generated EC-specific *Dhh* knockout mice (Cdh5^Cre^*Dhh*^fl/fl^; hereafter referred to as *Dhh*^ecKO^) (Fig. S5A). Successful deletion of *Dhh* was confirmed by genotyping (Fig. S5B) and RT-qPCR analysis (Fig. S5C). In *Dhh*^ecKO^ mice, *Dhh* mRNA expression was markedly reduced in the aortic intima (EC), whereas its expression in the medial layer (Non-EC) remained unchanged (Fig. S5C), supporting endothelial-specific deletion. Since DHH is a secreted protein, we additionally measured circulating DHH levels by ELISA and observed a significant reduction in serum DHH levels in *Dhh*^ecKO^ mice compared with control littermates (Fig. S5D). Importantly, endothelial *Dhh* deletion did not alter the expression of the other hedgehog ligands, *Shh* and *Ihh*, within the aortic intima (Fig. S2B and S5E), further supporting the specificity of the model. Consistently, IF staining of the brachiocephalic artery (BCA) demonstrated reduced DHH expression within the endothelial layer of *Dhh*^ecKO^ mice relative to controls (Fig. S5F). Collectively, these findings confirm efficient and endothelial-specific deletion of *Dhh* in *Dhh*^ecKO^ mice.

To investigate the role of endothelial-derived *Dhh* in DF-induced atherosclerosis, 8-week-old male and female *Dhh*^ecKO^ mice and their control littermates (*Dhh*^fl/fl^) received a single intravenous injection of adeno-associated virus 8 (AAV8) vector encoding *Pcsk9*-D377Y to promote hepatic low-density lipoprotein receptor (LDLR) degradation (Fig. S6) to induce hypercholesterolemia, and were maintained on a high-cholesterol diet (HCD) for 2 weeks before partial carotid ligation (PCL) surgery. Mice were subsequently maintained on HCD for an additional 3 weeks following PCL (Fig. 3A). No significant differences were observed between *Dhh*^ecKO^ and litter-mate control mice in body weight, blood glucose levels (Fig. S7A and S7D), serum lipid profiles, including total cholesterol (CHO), triglycerides (TG), low-density lipoprotein cholesterol (LDL), and high-density lipoprotein cholesterol (HDL) (Fig. S7B and S7E), or liver injury markers, including alanine aminotransferase (ALT) and aspartate aminotransferase (AST) (Fig. S7C and S7F).

**Fig. 3.**
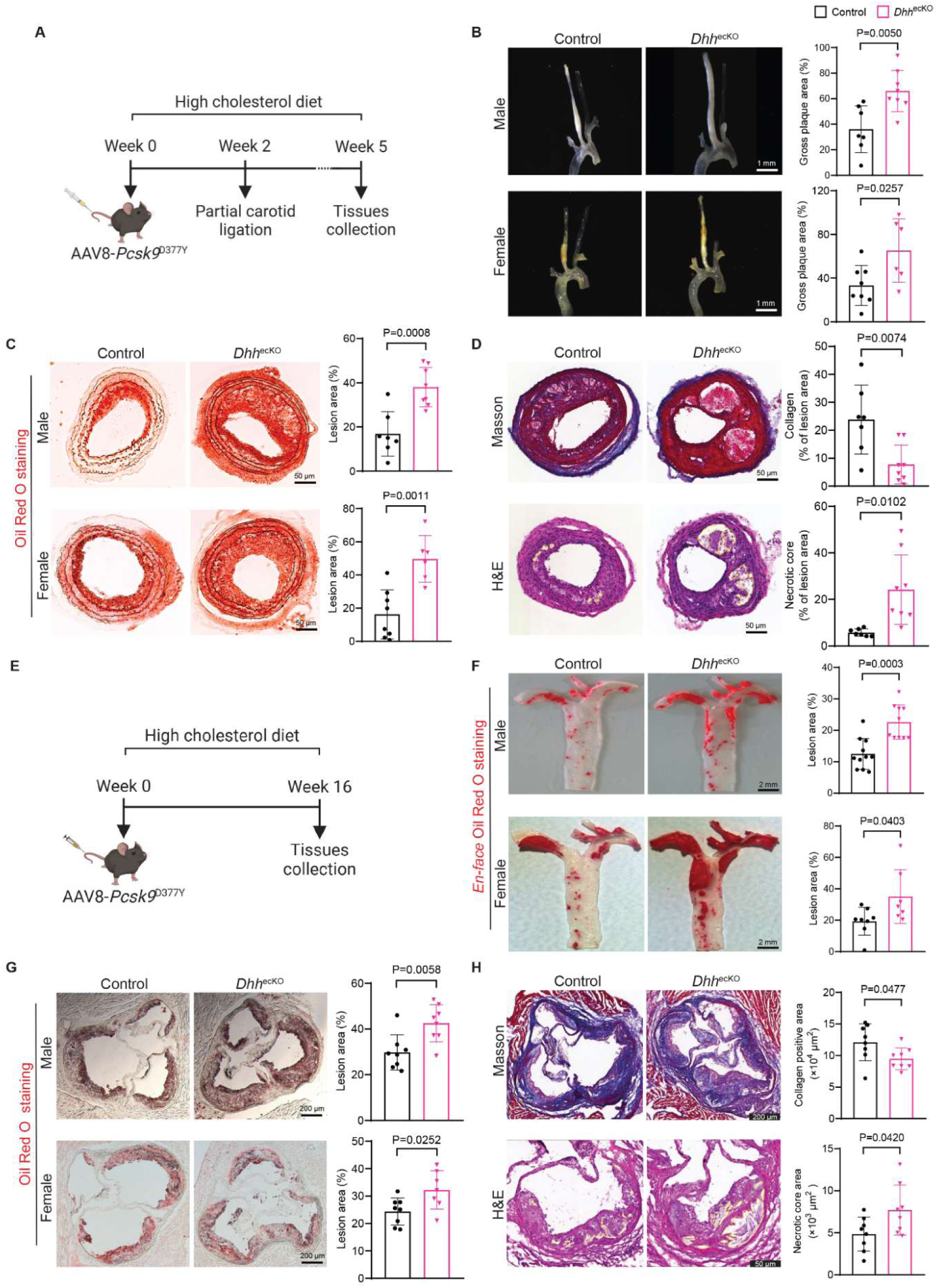
EC-specific deletion of *Dhh* exacerbates atherosclerosis development in mouse models. **(A**) Schematic diagram of experimental design. Control (*Dhh*^fl/fl^) and *Dhh*^ecKO^ (Cdh5^Cre^*Dhh*^fl/fl^) mice were intravenously given a single dose of AAV8-*Pcsk9*-D377Y for 2-weeks, followed by 3-week PCL surgery and HCD feeding. (**B)** Representative gross LCA images and lesion area quantification of the control and *Dhh*^ecKO^ male and female mice after 3-weeks of PCL (n=7 to 8 for males and n=8 to 6 for females). (**C)** Representative images and quantification data of Oil Red O staining of the control and *Dhh*^ecKO^ male and female mice LCA sections after 3-weeks of PCL (n=7 to 8 for males and n=8 to 6 for females). (**D)** Representative images and quantification data of Masson trichrome staining and H&E staining of the control and *Dhh*^ecKO^ male mice LCA sections after 3-weeks of PCL (n=7 to 8). (**E**) Schematic diagram of experimental design. Control and *Dhh*^ecKO^ mice were intravenously given a single dose of AAV8-*Pcsk9*-D377Y and maintained on HCD for 16-weeks. (**F**) Representative images and quantification data of *en-face* Oil Red O staining of the control and *Dhh*^ecKO^ male and female mice aorta after 16-weeks of AAV8-*Pcsk9* injection and HCD feeding (n=11 to 10 for males and n=8 to 7 for females). (**G)** Representative images and quantification data of Oil Red O staining of aortic sinus from the control and *Dhh*^ecKO^ male and female mice after 16 weeks of AAV8-*Pcsk9* injection and HCD feeding (n=8 to 8 for males and n=8 to 7 for females). (**H**) Representative images and quantification data of Masson trichrome staining and H&E staining of aortic sinus from the control and *Dhh*^ecKO^ male mice after 16 weeks of AAV8-*Pcsk9* injection and HCD feeding (n=8). All data are presented as mean ± SEM. Statistical analysis was performed using unpaired two-tailed t-test for **B**, **C**, **D**, **F**, **G** and **H**.

Interestingly, EC-specific deletion of *Dhh* significantly increased atherosclerotic lesion formation in the LCA of both male and female *Dhh*^ecKO^ mice following PCL (Fig. 3B). Consistent with this, Oil-Red O staining demonstrated increased lipid accumulation within lesions of *Dhh*^ecKO^ mice compared with controls (Fig. 3C). In addition, Masson’s trichrome staining revealed reduced collagen content within plaques, whereas hematoxylin and eosin (H&E) staining demonstrated enlarged necrotic core areas in the LCA of *Dhh*^ecKO^ mice (Fig. 3D). These findings suggest that endothelial *Dhh* deficiency promotes DF-associated atherosclerosis and is associated with features of plaque instability.

To further examine the role of endothelial *Dhh* to atherosclerosis progression under chronic hypercholesterolemic conditions, male and female *Dhh*^ecKO^ and control mice were intravenously injected with AAV8-*Pcsk9* and maintained on HCD for 16 weeks (Fig. 3E). Similar to the PCL model, no significant differences were observed in body weight, blood glucose levels (Fig. S8A and S8D), or serum lipid parameters, including CHO, TG, LDL, and HDL, between *Dhh*^ecKO^ and control mice (Fig. S8B and S8E). A modest increase in ALT and AST levels was observed in male *Dhh*^ecKO^ mice (Fig. S8C), whereas no differences were detected in female mice (Fig. S8F). In addition, male *Dhh*^ecKO^ mice exhibited increased diastolic blood pressure at 16 and 24 weeks of age, although systolic blood pressure did not differ significantly between groups (Fig. S9).

Notably, Oil Red O staining showed a marked increase in atherosclerotic plaque burden within both the whole aorta and aortic sinus of male and female *Dhh*^ecKO^ mice compared with controls (Fig. 3F and 3G). Consistent with the PCL model, plaques from *Dhh*^ecKO^ mice exhibited reduced collagen content and enlarged necrotic core areas, as assessed by Masson’s trichrome and H&E staining, respectively (Fig. 3H).

Together, these findings demonstrate that endothelial-specific deletion of *Dhh* accelerates atherosclerosis progression in hypercholesterolemic mice and is associated with features of unstable plaque remodeling.

### scRNA-seq analysis of *Dhh*^ecKO^ mice identifies *Dhh* as a novel regulator of EndoMT

To further define the endothelial-specific role of *Dhh* during atherosclerosis progression, we performed scRNA-seq analysis of aortic cells isolated from *Dhh*^ecKO^ and control mice following AAV8-*Pcsk9* administration and 16 weeks of HCD feeding (Fig. 4A). After preprocessing and quality-control filtering, 16,128 cells were retained for downstream analysis. Using the Seurat platform, we initially identified 18 distinct cell clusters, which were subsequently annotated into 6 major cell populations based on established marker genes for vascular and immune cell types within the mouse aorta, including macrophages, fibroblasts, VSMCs, ECs, T cells, and granulocytes (Fig. 4B and Fig. S10A-S10C).

**Fig. 4.**
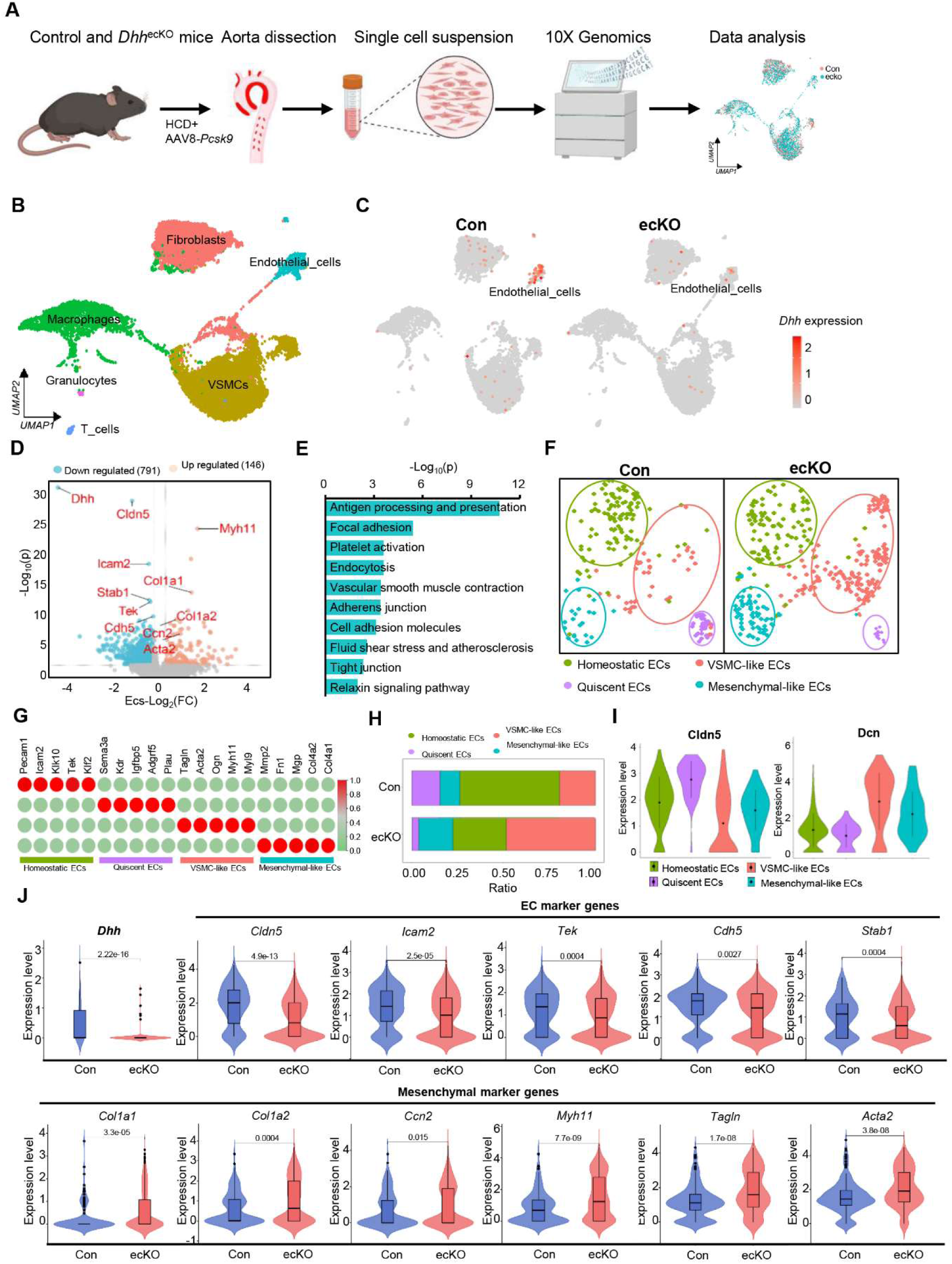
scRNA-seq analysis of *Dhh*^ecKO^ mice reveals *Dhh* as an endogenous suppressor of EndoMT. (**A**) Workflow of control and *Dhh*^ecKO^ scRNA-seq analysis. (**B**) UMAP plot exhibits the cell identification results from scRNA-seq analysis. (**C**) Feature Plot indicates specific knockout of *Dhh* in ECs (**D**) Volcano plot displays differentially expressed genes between control and *Dhh*^ecKO^ mice ECs. (**E**) KEGG signaling pathways for differentially expressed genes between groups of ECs. (**F**) UMAP plot displays the results of the sub-clustering of ECs. (**G**) Heatmap displays marker genes of each subpopulation of ECs. (**H**) Cell proportion chart shows the results of the proportion of each subpopulation of ECs. (**I**) Violin plot of representative marker genes for EC subpopulations. (**J**) Relative expression of marker genes for ECs and mesenchymal cells in the aorta from control and *Dhh*^ecKO^ mice. Statistical analysis was performed using the wilcox.test for **J**.

To assess the impact of endothelial *Dhh* deletion on hedgehog signaling, we examined the expression of key hedgehog pathway genes in ECs from control and *Dhh*^ecKO^ mice, as shown by heatmap in Fig. S2B. While *Dhh* expression was markedly reduced in ECs from *Dhh*^ecKO^ mice, no appreciable differences were observed in the expression of the other hedgehog ligands, *Shh* and *Ihh* (Fig. S2B), suggesting that endothelial *Dhh* deletion did not broadly alter hedgehog ligand expression within ECs. Feature plot analysis demonstrated a marked reduction in *Dhh* expression specifically within ECs from *Dhh*^ecKO^ mice (Fig. 4C), which was further confirmed by differential gene expression analysis shown in the volcano plot (Fig. 4D). To evaluate the specificity of endothelial *Dhh* deletion, we also examined *Dhh* expression in non-endothelial cell populations. *Dhh* expression remained unchanged in macrophages, other immune cell populations (Fig. S11A), and isolated peripheral blood mononuclear cells (PBMCs) from *Dhh*^ecKO^ mice compared with controls (Fig. S11B), supporting the endothelial specificity of the knockout model.

Differential expression analysis further revealed substantial alterations in endothelial-identity genes, including *Cldn5, Icam2, Stab1, Tek,* and *Cdh5*, as well as genes associated with mesenchymal and smooth muscle phenotypes, including *Myh11, Col1a1, Col1a2, Ccn2* and *Acta2* (Fig. 4D). KEGG pathway enrichment analysis of differentially expressed genes in ECs identified pathways related to focal adhesion, platelet activation, endocytosis, vascular smooth muscle contraction, adherens junctions, cell adhesion molecules, and fluid shear stress and atherosclerosis (Fig. 4E), suggesting that endothelial *Dhh* may influence EC phenotypic regulation through these signaling networks.

To further characterize endothelial phenotypic changes associated with *Dhh* deletion, we performed subclustering analysis of EC populations identified by scRNA-seq. Based on the expression of characteristic marker genes, ECs were classified into 4 subpopulations: homeostatic ECs, quiescent ECs, VSMC-like ECs, and mesenchymal-like ECs (Fig. 4F). The expression profiles of representative marker genes for each EC subset are shown in the heatmaps in Fig. 4G and Fig. S10D. Analysis of EC subset composition demonstrated that VSMC-like and mesenchymal-like EC populations were increased in *Dhh*^ecKO^ mice, whereas homeostatic and quiescent EC populations were reduced (Fig. 4H). Consistent with these changes, violin plot analysis showed reduced expression of endothelial homeostatic markers, including *Cldn5, Icam2, Tek, Cdh5,* and *Stab1*, in *Dhh*^ecKO^ ECs, whereas genes associated with mesenchymal and smooth muscle phenotypes, including *Dcn, Col1a1, Col1a2, Ccn2, Myh11, Tagln,* and *Acta2*, were increased (Fig. 4I and 4J). These transcriptional alterations were further validated by RT-qPCR analysis of endothelial-enriched intimal samples isolated from control and *Dhh*^ecKO^ mice (Fig. S12).

Collectively, these findings suggest that endothelial *Dhh* contributes to the maintenance of endothelial homeostasis and that its deletion promotes a phenotypic shift toward VSMC-like and mesenchymal-like endothelial states during atherosclerosis progression.

### Endothelial *DHH* inhibits EndoMT

Because endothelial-specific deletion of *Dhh* promoted EndoMT-associated phenotypic changes in mouse ECs, we next investigated whether *DHH* depletion induces a similar response in human ECs. To this end, we performed bulk RNA sequencing of *DHH*-silenced HUVECs (schematic shown in Fig. 5A). Transcriptomic analysis revealed marked downregulation of endothelial-associated genes, including *KLF2, MEF2A, THBD, PLAU, GCH1,* and *NOS3,* together with increased expression of genes associated with EndoMT and mesenchymal phenotypes, including *POSTN, TAGLN, FAP, SNAI1, COL1A2, TGFBR1, SERPINE1, CCN2,* and *ZEB2* (Fig. 5B). Efficient *DHH* knockdown and the expression changes of representative endothelial and EndoMT-related genes are shown in Fig. 5C and 5D.

**Fig. 5:**
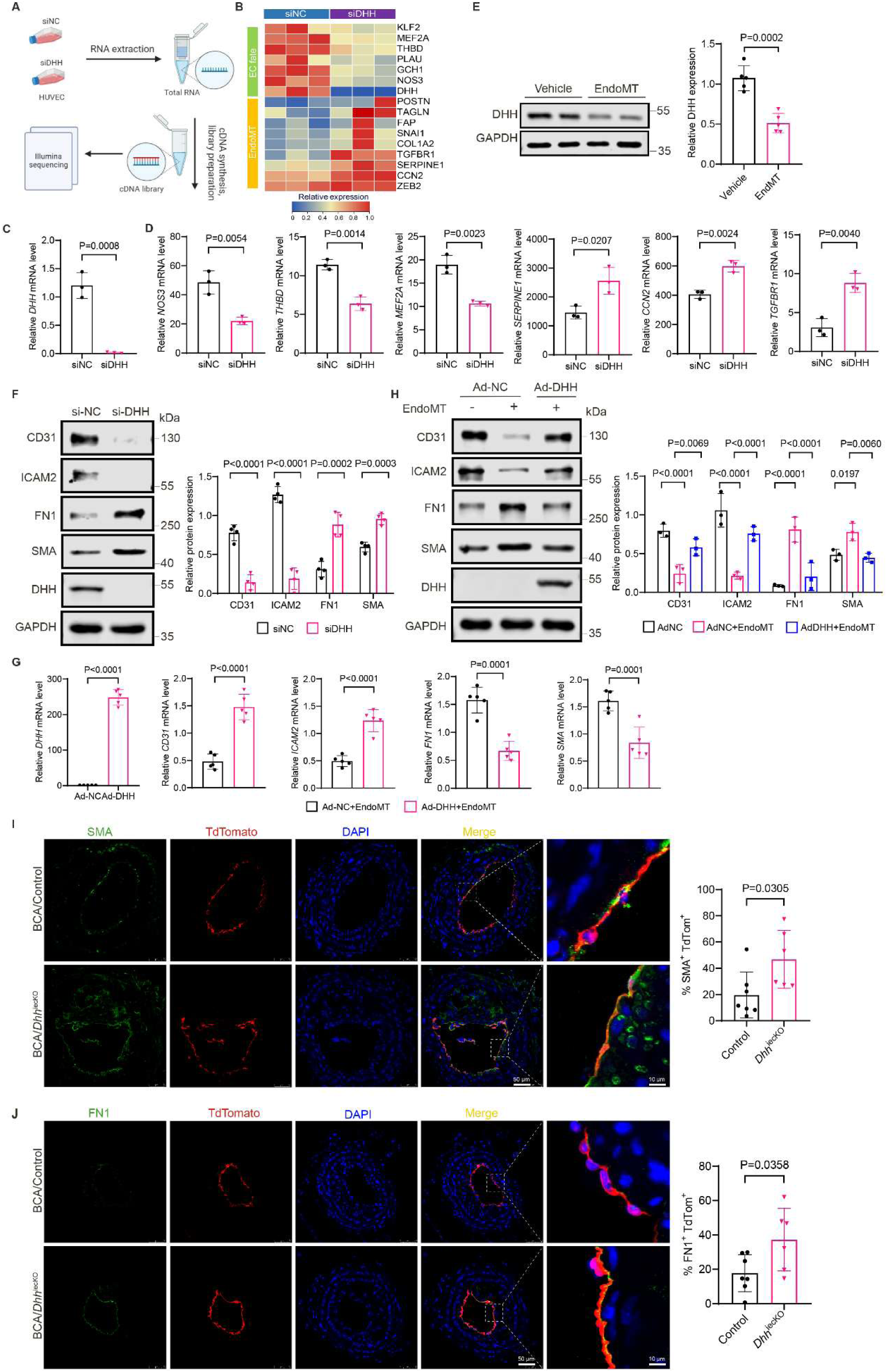
Endothelial *DHH* inhibits EndoMT *in vitro* and *in vivo*. **(A**) Workflow of *DHH* de-pleted HUVECs RNA-sequencing. (**B**) Heat map analysis of the EC marker genes and EndoMT marker genes of RNA-sequencing HUVECs. (**C**) Relative expression of *DHH* in siNC and siDHH-treated HUVECs derived from bulk RNA-sequencing analysis (n=3). (**D**) Relative ex-pression of *NOS3, THBD, MEF2A, SERPINE1, CCN2* and *TGFBR1* in siNC and siDHH-treated HUVECs derived from bulk RNA-sequencing analysis (n=3). (**E**) Representative Western blots and quantification of DHH protein in HAECs treated with vehicle or EndoMT triggers (hTGF-β1+hIL-1β+hTNF-α, each cytokine concentration is 10 ng/mL) for 48 h (n=5). (**F**) Representa-tive Western blots and quantification of CD31, ICAM2, FN1 and SMA in HAECs transfected with siNC or siDHH for 3 days (n=3). (**G**) RT-qPCR analysis of mRNA levels of *DHH*, *CD31*, *ICAM2*, *FN1,* and *SMA* in HAECs transfected with Ad-NC or Ad-DHH and treated with En-doMT triggers for 48 h (n=5). (**H**) Representative Western blots and quantification of CD31, ICAM2, FN1 and SMA in HAECs transfected with Ad-NC or Ad-DHH and treated with vehicle or EndoMT triggers for 3 days (n=3). (**I** and **J)** Representative immunofluorescence staining im-ages of brachiocephalic artery (BCA) sections from control and *Dhh*^iecKO^ mice stained for TdTomato^+^ (red), SMA^+^ (green) and FN1^+^ (green). Quantification was performed by calculating the percentage of SMA^+^ TdTomato^+^ or FN1^+^ TdTomato^+^ copositive cells (n=7 to 6). All data are presented as mean ± SEM. Statistical analysis was performed using unpaired two-tailed Student t-test for **C, D, E, G, I** and **J.** One-way ANOVA followed by the Bonferroni method is used for **F.** Two-way ANOVA followed by the Bonferroni method of multiple pairwise comparisons test is used for **H.**

To further examine the functional role of *DHH* in EndoMT *in vitro*, we evaluated the expression of endothelial and mesenchymal markers in HAECs (human aortic endothelial cells) upon altered *DHH* levels. EndoMT was induced using a cytokine cocktail containing TGF-β1, IL-1β, and TNF-α, which are established inducers of endothelial phenotypic transition.^20, 27^ Cytokine treatment significantly reduced DHH protein expression in HAECs (Fig. 5E), suggesting an association between *DHH* downregulation and EndoMT induction. Consistent with this observation, *DHH* knockdown significantly reduced the expression of endothelial markers, including CD31 and ICAM2, while increased the mesenchymal markers FN1 and SMA at the protein level (Fig. 5F), indicating induction of EndoMT. In contrast, DHH overexpression attenuated cytokine-induced EndoMT-associated changes by restoring CD31 and ICAM2 expression and suppressing FN1 and SMA expression at both the mRNA and protein levels (Fig. 5G and 5H). These findings suggest that endothelial *DHH* negatively regulates EndoMT *in vitro*.

We next investigated whether *DHH*-mediated regulation of EndoMT involves canonical hedgehog signaling. Analysis of intimal ECs isolated from *Dhh*^ecKO^ and control mice revealed no significant changes in the expression of canonical hedgehog downstream genes, including *Ptch1, Ptch2, Gli1, Gli2,* and *Gli3* (Fig. S13A). To confirm inhibition of canonical hedgehog signaling, we performed a Gli-responsive dual-luciferase reporter assay, which demonstrated that sonidegib effectively suppressed Gli1 transcriptional activity (Fig. S13B). However, sonidegib treatment neither altered *DHH* knockdown-induced EndoMT (Fig. S13C) nor reversed the inhibitory effects of *DHH* overexpression on EndoMT (Fig. S13D). Together, these findings suggest that the anti-EndoMT effects of *DHH* are probably independent of canonical hedgehog signaling and may involve alternative signaling mechanisms.

To determine whether endothelial *DHH* suppresses EndoMT *in vivo*, we generated inducible endothelial lineage-tracing mice by crossing Cdh5-CreERT2; Rosa26-tdTomato mice with *Dhh*^fl/fl^ mice (*Dhh*^iecKO^ lineage-tracing mice). Mice were subsequently administered tamoxifen for 5 consecutive days to induce Cre recombination and tdTomato expression. Following a 2-week washout period, mice received AAV8-*Pcsk9* and were maintained on a high-cholesterol diet for 9 weeks to induce atherosclerosis (Fig. S14A). Efficient deletion of *Dhh* was confirmed by genotyping (Fig. S14B) and RT-qPCR analysis (Fig. S14C). *Dhh* expression was markedly reduced in the aortic intima, whereas its expression in the medial layer remained unchanged, supporting endothe-lial-specific deletion (Fig. S14C). Consistently, immunofluorescence staining of the BCA demonstrated reduced DHH protein expression within the endothelial layer of *Dhh*^iecKO^ mice compared with controls (Fig. S14D). To validate endothelial-specific tdTomato labeling, immunostaining demonstrated colocalization of tdTomato with CD31 within the endothelial layer (Fig. S14E). No significant differences were observed in body weight, blood glucose levels, serum lipid profiles, or ALT and AST levels between *Dhh*^iecKO^ and control mice (Fig. S15A to S15C). Consistent with findings from the constitutive knockout and PCL models, inducible endothelial-specific deletion of *Dhh* significantly increased lesion formation in the BCA compared with controls (Fig. S15D). Importantly, immunofluorescence analysis demonstrated that a substantial proportion of FN1^+^ and SMA^+^ cells co-localized with tdTomato, confirming their endothelial origin (Fig. 5I and 5J). More-over, *Dhh*^iecKO^ mice exhibited a marked increase in tdTomato^+^SMA^+^ and tdTomato^+^FN1^+^ cells within atherosclerotic lesions of the BCA, indicating enhanced EndoMT following endothelial *Dhh* deletion (Fig. 5I and 5J).

Collectively, these findings provide both *in vitro* and *in vivo* evidence that endothelial *DHH* suppresses EndoMT under atherogenic conditions.

### DHH interacts with PAI-1 and suppresses PAI-1 dependent LRP1/AKT/ERK1/2 and LRP1/SMAD2/3 pathway

Next, we explored the direct mechanism by which *DHH* mitigates the EndoMT process. We precipitated DHH from overexpressed HUVECs and subjected to mass spectrometry (Fig. 6A). Among the identified proteins, PAI-1 (gene name: *SERPINE1*) had the highest unique peptide number and coverage (Fig. 6B) and has been reported to be closely associated with EndoMT.^24, 41^ Co-immunoprecipitation assays confirmed the interaction of DHH and PAI-1 under EndoMT conditions in HUVECs (Fig. 6C). Molecular docking of DHH and PAI-1 protein revealed a high binding score of -311.68 kcal/mol, indicating a strong interaction between the two proteins (Fig. 6D). Our further analysis showed that amino acid residues A63, R66, Q137, N190, R195, E348, L394 of DHH protein and amino acid residues R187, A203, E242, K243, E244, E348, E392 of PAI-1 protein exhibited strong hydrogen bonding, confirming the direct binding between two proteins (Fig. 6E). Moreover, surface plasmon resonance (SPR) analysis showed that PAI-1 directly binds to the DHH protein in a concentration-dependent manner (Fig. 6F). Affinity fitting calculations yielded a preliminary KD value of 4.224^E-7^ M for the DHH and PAI-1 proteins interaction (Fig. 6G), suggesting that both proteins directly interact with each other.

**Fig. 6.**
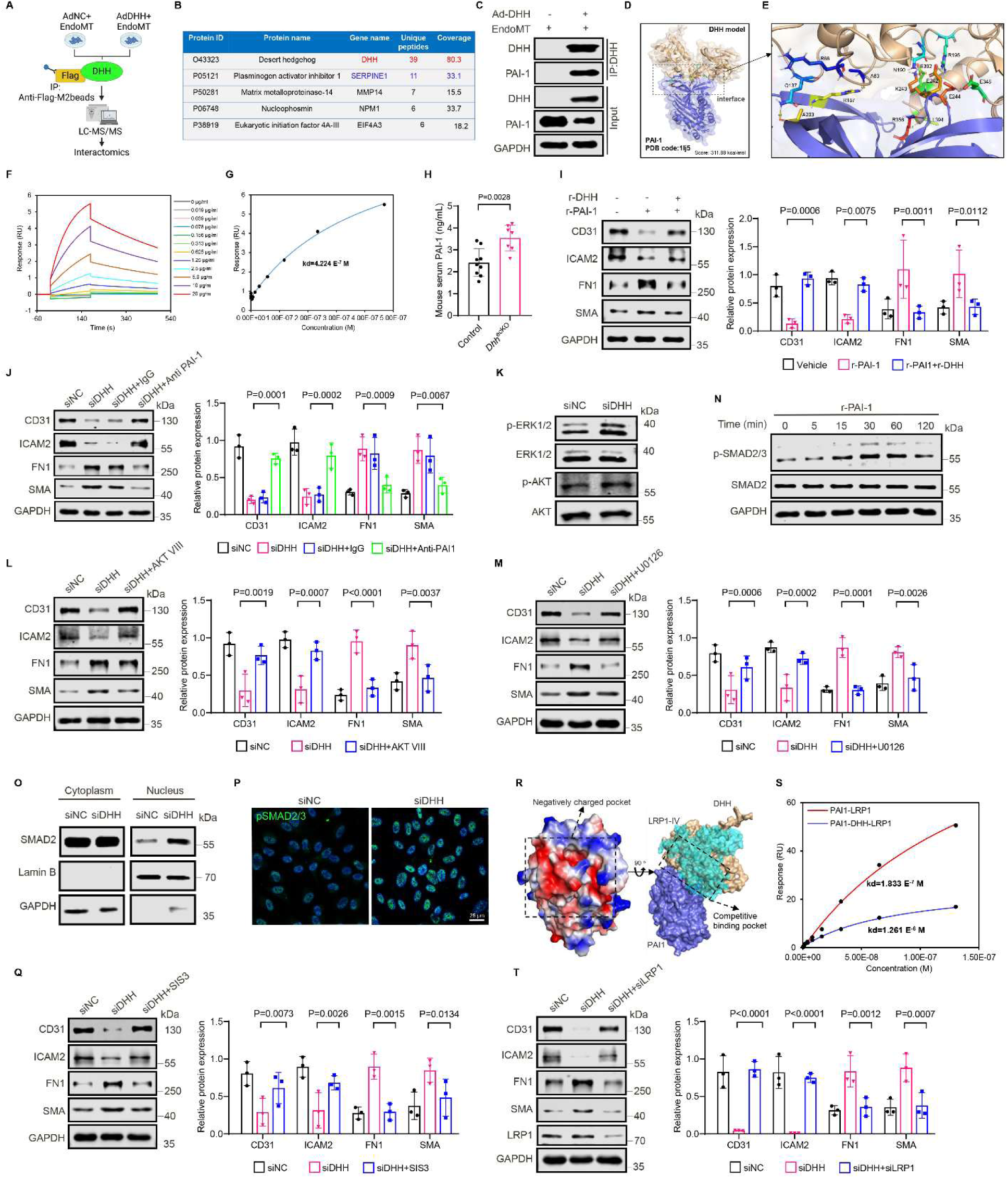
DHH interacts with PAI-1 and mitigates PAI-1 mediated LRP1/ AKT/ERK1/2 and LRP1/SMAD2/3 pathway. **(A**) Workflow of mass spectrometry of HUVECs transfected with Ad-NC or Ad-DHH and treated with EndoMT triggers for 3 days. Lysates were immunoprecipitated with anti-Flag-M2 beads. (**B**) Representative DHH interacting proteins identified by mass spectrometry analysis. (**C**) IP and immunoblot analysis of the DHH interaction with PAI-1 in HUVECs transfected with Ad-NC or Ad-DHH and treated with EndoMT triggers for 3 days. (**D**) Molecular docking of DHH and PAI-1 protein, with a docking score of -311.68 kcal/mol. (**E**) Cartoon representation showing strong hydrogen bonding between the amino acid residues A63, R66, Q137, N190, R195, E348, L394 of DHH protein and the amino acid residues R187, A203, E242, K243, E244, E348, E392 of PAI-1 protein. (**F**) Surface plasmon resonance (SPR) sensor-gram showing the binding kinetics of the interaction between DHH and PAI-1 proteins at various concentrations of PAI-1 (0-20 µg/ml). The colored curves represent different PAI-1 protein concentrations, with responses recorded in real time (Resonance Units, RU). (**G**) 4.224 E^-7^ M kd value for DHH and PAI-1 calculated based on SPR response values, indicating strong binding between the two proteins. (**H)** The concentration of PAI-1 in the control and *Dhh*^ecKO^ mice serum was detected by ELISA (n=9 to 7). (**I**) Representative Western blots and quantification of CD31, ICAM2, FN1 and SMA in HAECs treated with r-DHH (100 ng/mL) or r-PAI-1 (100ng/mL) for 3 days (n=3). (**J**) Representative Western blots and quantification of CD31, ICAM2, FN1 and SMA in HAECs transfected with siNC or siDHH for 24 h, followed by treatment with control anti-IgG (10 µg/mL) or PAI-1 neutralizing antibody (10 µg/mL) for an additional 24 h (n=3). (**K**) Representative Western blots of p-ERK1/2, ERK1/2, p-AKT and AKT in HUVECs transfected with siNC or siDHH for 48 h (n=3). **(L)** Representative Western blots and quantification of CD31, ICAM2, FN1 and SMA in HAECs transfected with siNC or siDHH for 24 h, followed by treatment with AKT inhibitor VIII (300 nM) for an additional 24 h (n=3). (**M)** Representative Western blots and quantification of CD31, ICAM2, FN1 and SMA in HAECs transfected with siNC or siDHH for 24 h, followed by treatment with ERK1/2 inhibitor U0126 (60 nM) for an additional 24 h (n=3). (**N**) Representative Western blots of p-SMAD2/3, SMAD2 and GAPDH in HUVECs treated with rPAI-1 protein (100ng/mL) for different time points (n=6). (**O**) Representative Western blots of cytoplasm and nuclear fractions of SMAD2, Lamin B and GAPDH in HUVECs transfected with siNC or siDHH for 48 h (n=3). (**P**) Representative immunostaining images of pSMAD2/3 (green) and DAPI (blue) in the HAECs transfected with siNC or siDHH for 48 h (n=3). **(Q)** Representative Western blots and quantification of CD31, ICAM2, FN1 and SMA in HAECs transfected with siNC or siDHH for 24 h, followed by treatment with SMAD3 inhibitor SIS3 (3 µM) for an additional 24 h (n=3). (**R**) The protein electrostatic potential map and cartoon diagram reveal that the PAI-1 protein possesses a distinct negative charge pocket, and both DHH and LRP1 cluster IV bind to this negative charge pocket of PAI-1. (**S**) SPR measurements indicate that the kd between PAI-1 and LRP1 cluster IV is 1.833 E^-7^ M. However, when PAI-1 binds to DHH, the kd for the binding of LRP1 cluster IV to PAI-1 increases significantly to 1.261 E^-6^ M. (**T**) Representative Western blots and quantification of CD31, ICAM2, FN1 and SMA in HAECs transfected with siNC or siLRP1 for 24 h, followed by transfection with siDHH for an additional 48 h (n=3). All data are presented as mean ± SEM. Statistical analfysis was performed using unpaired two-tailed t-test for **H** and two-way ANOVA followed by the Bonferroni method of multiple pairwise comparisons test for **I, J, L, M, Q** and **T.**

Intriguingly, *DHH* knockdown in HUVECs increased PAI-1 protein expression (Fig. S16A). While, *DHH* overexpression reduced PAI-1 expression in a dose-dependent manner (Fig. S16B). Consistently, circulating PAI-1 levels were significantly elevated in the serum of *Dhh*^ecKO^ mice compared to control mice (Fig. 6H). To determine whether *DHH* regulates EndoMT and atherosclerosis through PAI-1, we performed functional studies. Treatment of HAECs with recombinant human PAI-1 (hrPAI) protein induced EndoMT, whereas co-treatment with recombinant human DHH (hrDHH) protein reversed this effect (Fig. 6I). Furthermore, immunodepletion of secreted PAI-1 from conditioned media of DHH knockdown HAECs using PAI-1-neutralizing antibody significantly abolished *DHH* knockdown-induced EndoMT phenotype, as assessed by Western blot analysis (Fig. 6J).

Next, to explore how DHH prevents PAI-1-induced EndoMT, we first examined the involvement of the LRP1/AKT/ERK1/2 signaling pathway, as PAI-1 has been reported to induce EndoMT through LRP1-dependent AKT and ERK1/2 phosphorylation in lymphatic endothelial cells.^24, 42^ *DHH* knockdown induced phosphorylation of ERK1/2 and AKT, closely resembling the effects of PAI-1 (Fig. 6K). Furthermore, treatment of HUVECs with AKT or ERK1/2 inhibitors abolished *DHH* knockdown-induced EndoMT (Fig. 6L and 6M). Since the TGF-β pathway is a critical driver of EndoMT, we next assessed its involvement in *DHH* deficiency-induced EndoMT. We found that PAI-1 activated TGF-β signaling by inducing Smad2/3 phosphorylation (Fig. 6N), similar to its effects on AKT/ERK1/2. Furthermore, our western blot analysis showed that *DHH* knockdown significantly increased Smad2 translocation to the nucleus (Fig. 6O). IF staining further verified increased Smad2 translocation in *DHH* knockdown HAECs (Fig. 6P), suggesting DHH’s depletion role in Smad2/3 pathway activation. Interestingly, treatment of HAECs with Smad3 inhibitor SIS3 prevented *DHH* knockdown-induced EndoMT (Fig. 6Q).

Finally, we investigated whether these signaling events are dependent on PAI-1 receptor, LRP1. The molecular docking results revealed that both DHH and LRP1 cluster IV bind to the negatively charged pocket of PAI-1, exhibiting a competitive relationship (Fig. 6R). SPR analysis confirmed that LRP1 cluster IV binds strongly to PAI-1 (Kd = 1.833 × 10⁻⁷ M), but the binding affinity significantly decreased when PAI-1 binds to DHH (Kd = 1.261 × 10⁻⁶ M) (Fig. 6S). Additionally, siRNA-mediated knockdown of LRP1 abolished DHH knockdown-induced EndoMT (Fig. T). These results indicate that DHH competes with LRP1 to block the PAI-1-LRP1 interaction and retards PAI-1-induced EndoMT by inhibiting the PAI-1-dependent LRP1/AKT/ERK1/2 and LRP1/SMAD2/3 signaling pathways.

To extend these findings *in vivo*, we performed pharmacological inhibition studies using the selective PAI-1 inhibitor TM5275.^43^ 8-week-old male *Dhh*^ecKO^ mice and their control littermates received a single intravenous injection of AAV8-*Pcsk9* and were maintained on HCD for 2 weeks before PCL surgery. TM5275 was administered orally at a dose of 20 mg/kg/day, beginning 1 week prior to PCL surgery and continuing for a total of 4 weeks (experimental schematic shown in Fig. S17A). Notably, TM5275 treatment significantly reduced atherosclerotic lesion formation in the LCA of *Dhh*^ecKO^ mice, as assessed by gross images of the LCA as well as Oil-Red O staining of LCA sections (Fig. S17B and S17C). No significant differences were observed in the body weight, blood glucose levels and serum lipid profiles, among control, *Dhh*^ecKO^ and TM5275 treated *Dhh*^ecKO^ group (Fig. S17D and S17E). Collectively, these findings suggest that PAI-1 functions downstream of DHH signaling in the regulation of EndoMT and atherosclerosis progression.

### Recombinant DHH Prevents EndoMT and Atherosclerosis

To assess the *in vivo* effects of recombinant DHH (rDHH) protein on EndoMT and atherosclerosis, we used *ApoE* KO mice. After 4-weeks of HCD feeding to establish atherosclerotic plaques, *ApoE* KO mice were treated with N-terminal domain of rDHH protein (2 mg/kg*, i.p.*) or vehicle three times a week for 12-weeks concurrently HCD feeding (Fig. 7A and 7B). Notably, no phenotypic differences were observed in body weight, blood glucose levels, serum lipid levels, and ALT or AST levels between rDHH-treated mice compared to control mice (Fig. S18A to S18C). Interestingly, Oil Red O staining revealed a significant reduction in atherosclerotic plaque area within the aorta (Fig. 7C) and aortic sinus (Fig. 7D) of rDHH-treated *ApoE* KO mice compared to control mice. Masson’s trichrome staining showed more collagen deposition in rDHH-treated mice compared to their respective controls (Fig. 7E). H&E staining of the aortic sinus revealed smaller necrotic core areas in rDHH-treated mice (Fig 7F), indicating the features of stable plaques.

**Fig. 7.**
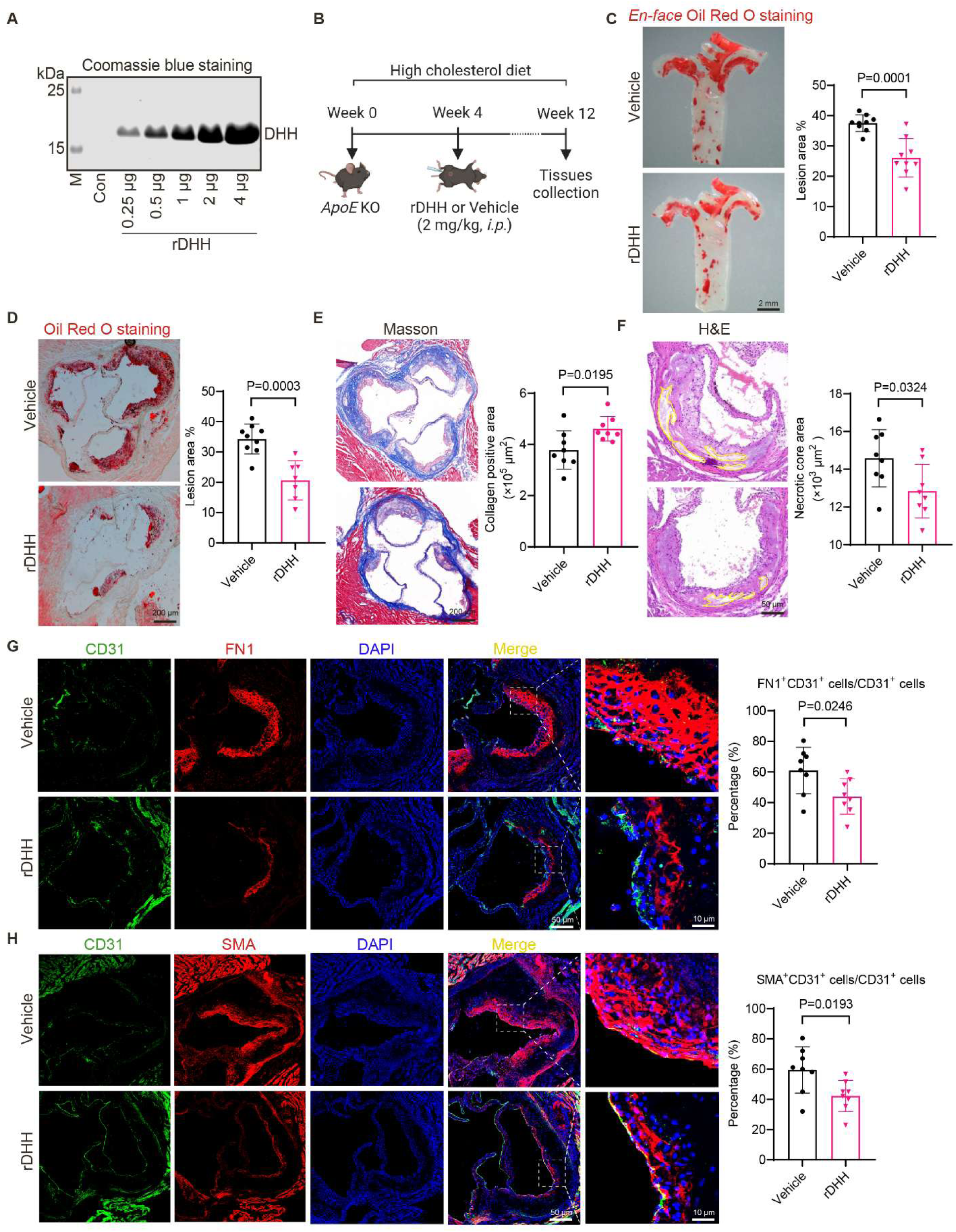
Recombinant DHH protein prevents EndoMT and atherosclerosis. **(A**) Recombinant mouse DHH (rDHH) protein purification and expression validation by Coomassie blue staining. (**B**) Schematic diagram of experimental design. *ApoE* KO mice were fed a high cholesterol diet and treated with rDHH protein (2 mg/kg, *i.p.*) or vehicle three times a week for 8weeks. (**C**) Representative images and quantification data of *enface* Oil Red O staining of the vehicle and rDHH treated *ApoE* KO mice aorta after 12-weeks of HCD feeding and 8-weeks of vehicle (PBS) or rDHH protein treatment (n=9). (**D** to **F**) Representative images and quantification data of Oil Red O staining (**D)**, Masson trichrome staining (**E),** and H&E staining (**F)** of the aortic sinus of the vehicle and rDHH treated *ApoE* KO mice (n=8). (**G and H**) Representative immuno-fluorescence staining images of the aortic sinus of the vehicle and rDHH-treated *ApoE* KO mice stained for CD31 (green), FN1 (red) and SMA (red) (n=8). Quantification was performed by dividing FN1^+^CD31^+^ or SMA^+^CD31^+^ copositive cells over total CD31^+^ cells. All data are presented as mean ± SEM. Statistical analysis was performed using unpaired two-tailed Student t-test for **C**, **D**, **E**, **F, G** and **H**.

Importantly, rDHH treatment significantly inhibited EndoMT *in vivo,* as evidenced by the increase in CD31^+^ ECs number and a substantial decrease in CD31^+^FN1^+^ or CD31^+^SMA^+^ copositive cells in the aortic sinus of rDHH treated group as compared to controls (Fig. 7G and 7H). These results indicate that recombinant DHH protein treatment prevents EndoMT and atherosclerosis development in *ApoE* KO mice and favors a stable plaque phenotype.

### DHH negatively correlates with PAI-1 expression in patients with CAD

Finally, we investigated whether DHH expression is associated with atherosclerotic plaque severity in human disease. Analysis of a publicly available dataset (GSE149759) containing stable and unstable plaques from individuals with coronary artery disease (CAD) revealed that *DHH* mRNA expression was significantly reduced in unstable plaques compared with stable plaques (Fig. 8A). In contrast, *SERPINE1* mRNA expression was increased in unstable plaques (Fig. 8A). To further evaluate this association in an independent clinical cohort, we analyzed gene expression profiles from advanced carotid atherosclerotic plaques obtained from the Athero-Express Biobank, an on-going prospective cohort of patients undergoing carotid endarterectomy at the University Medical Center Utrecht and St. Antonius Hospital, Nieuwegein, the Netherlands.^44^ Among 981 symptomatic carotid endarterectomy patients, plaque *DHH* expression negatively correlated with *SERPINE1* expression, supporting an inverse relationship between *DHH* and *SERPINE1* expression in advanced human atherosclerosis (Fig. 8B).

**Fig. 8.**
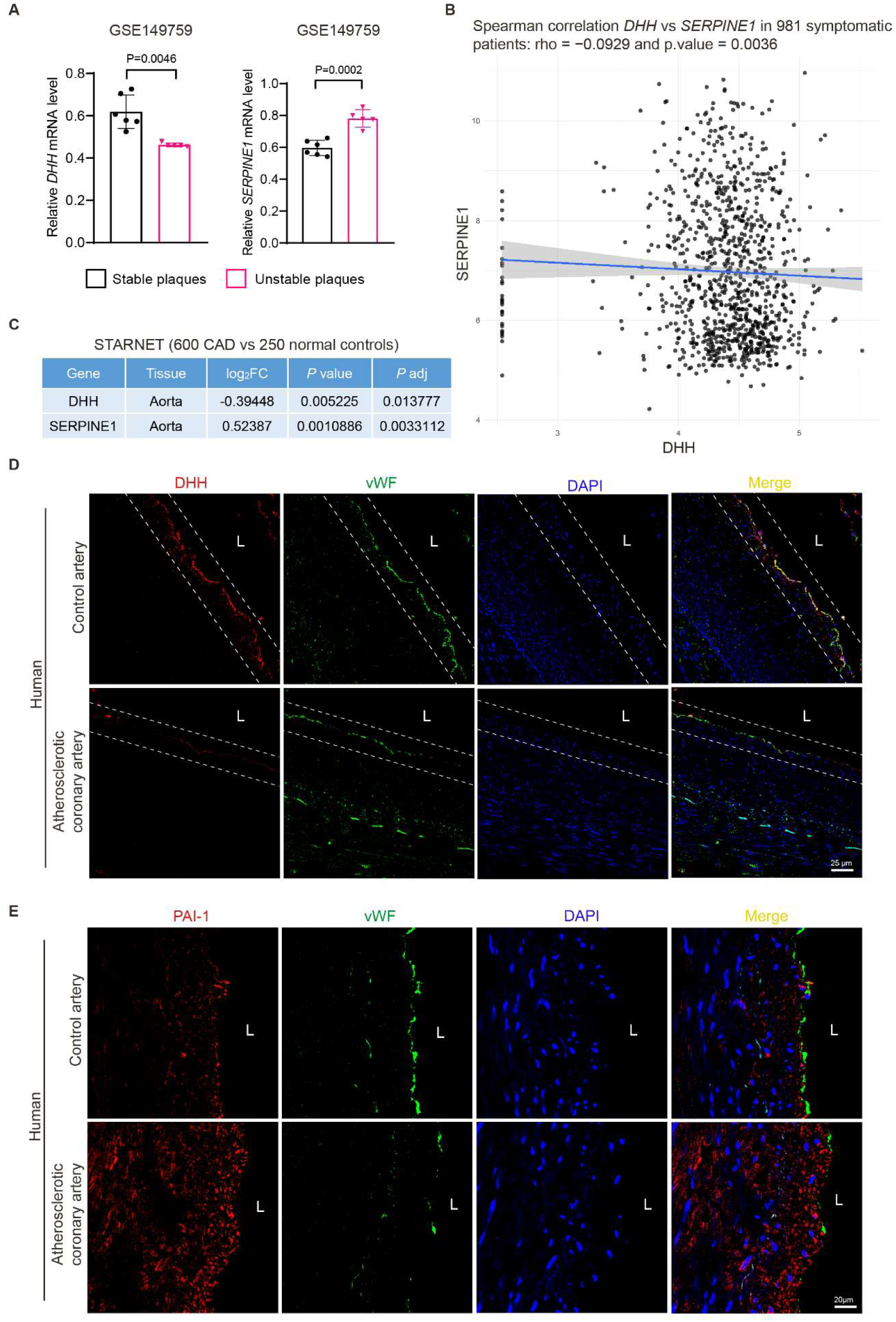
DHH negatively correlates with PAI-1 expression in patients with CAD. **(A**) The gene expression of *DHH* and *SERPINE1* (PAI-1) in the stable and unstable plaques of CAD patients in a public dataset (GSE149759) (n=6 to 5). (**B**) *DHH* and *SERPINE1* expression in plaque, within 981 patients who experienced symptoms before carotid endarterectomy. (**C**) *DHH* and *SERPINE1* expression in the aorta of CAD patients (n=600) and healthy controls (n=250) in the STARNET database. (**D**) Representative immunostaining images of DHH (red), vWF (green) and DAPI (blue) in the human control coronary arteries and atherosclerotic coronary arteries (n=3). (**E**) Representative immunostaining images of PAI-1 (red), vWF (green) and DAPI (blue) in the human control coronary arteries and atherosclerotic coronary arteries (n=3). All data are presented as mean ± SEM. Statistical analysis was performed using unpaired two-tailed t-test for **A** and Spearman’s rank correlation coefficient for **B**.

We next assessed DHH and PAI-1 expression in human aortic tissues using data from the STARNET (Stockholm-Tartu Atherosclerosis Reverse Network Engineering Task) database, which included samples from 600 patients with CAD and 250 healthy controls.^45, 46^ *DHH* expression was significantly decreased, whereas PAI-1 expression was increased in atherosclerotic aortic tissues from CAD patients compared with healthy controls (Figure 8C). To extend these findings at the protein level, we examined DHH and PAI-1 expression in coronary artery specimens from control individuals and patients with CAD. DHH expression was reduced, whereas PAI-1 expression was increased in atherosclerotic coronary arteries compared with non-atherosclerotic control arteries (Fig. 8D and 8E). Consistent with these findings, serum DHH levels were also lower in patients with CAD than in healthy individuals (Fig. S19). Collectively, these data support an association between reduced DHH expression and increased PAI-1 expression in human atherosclerotic disease.

## Discussion

DHH is a hedgehog ligand that shows preferential expression in ECs, with its biological functions in vascular biology and diseases being largely unknown. Previously, a genome-wide microarray study using a mouse carotid artery ligation model has revealed *Dhh* downregulation in response to DF in arterial ECs *in vivo.*^40^ However, the precise role of *DHH* in atherosclerosis remains elusive. Given the nature of atherosclerosis as a biomechanical disease with focal development of plaques in the regions of DF, the mechanoresponsive feature of *DHH* may implicate *DHH* in regulating atherosclerosis. Our scRNA-seq study lends support to the downregulation of *Dhh* in the atheroprone AA compared with atheroresistant descending TA in the *ApoE* KO mice. The salient findings of this study can be summarized as follows: (1) EC-specific deletion of *Dhh* exacerbates atherosclerotic plaque development in well-established murine models of atherosclerosis. (2) Endothelial loss of *Dhh* promotes EndoMT. (3) DHH interacts with PAI-1 and retards PAI-1-dependent EndoMT via suppressing PAI-1 binding to LRP1 and ensuing AKT/ERK1/2 and SMAD2/3 pathways. (4) The recombinant mouse DHH protein administration inhibits EndoMT and atherosclerosis development in *ApoE* KO mice. (5) Human coronary arteries with advanced plaques and serum from CAD patients showed decreased endothelial DHH expression/secretion. These findings coherently pinpoint an essential role of DHH in maintaining endothelial resilience in the face of pro-atherogenic risk factors.

Previously, *DHH* knockdown in ECs was reported to promote the expression of adhesion molecules such as VCAM-1, ICAM-1 and other inflammatory cytokines, including IL-6 and CCL2.^37^ Therefore, we first investigated whether *DHH* prevents atherosclerosis by inhibiting ECs inflammation. Interestingly, our results demonstrated that *DHH* does not seem to be involved in endothelial inflammation since *DHH* silencing or overexpression did not influence TNF-α and IL-1β induced protein expression of VCAM-1 or ICAM-1 (Fig. S20A and S20B). Additionally, *DHH* silencing or overexpression did not impact TNF-α-induced monocyte adhesion to HAECs (Fig. S20C and S20D). Consequently, we explored the potential mechanisms by which the *DHH* confers protection against atherosclerosis. Our unbiased scRNA-seq analysis revealed significant enrichment in VSMC-like/mesenchymal-like EC clusters and a substantial decrease in homeostatic/quiescent EC clusters in the aortic endothelium of *Dhh*^ecKO^ mice, suggesting that endothelial loss of *Dhh* dictates ECs transition from a quiescent state to a mesenchymal-like state. This phenotype was recapitulated in human ECs depleted of DHH.

Mechanistically, we identified PAI-1 as a potential DHH-interacting protein. PAI-1 is a multifunctional secreted glycoprotein that serves as the primary physiological inhibitor of tissue-type (tPA) and urokinase-type (uPA) plasminogen activators. Apart from being crucially involved in fibrinolysis and wound healing, PAI-1 plays a pivotal role in cardiovascular disease, including myocardial infarction,^47^ ischemic stroke,^48^ coronary heart disease,^49^ and atherosclerosis.^50–52^ Ele-vated levels of PAI-1 downregulate tPA and uPA activity and create a prothrombotic or hypofibrinolytic state that contributes to cardiovascular disease pathogenesis.^52, 53^ Recently, PAI-1 has also been directly linked to EndoMT. Wei et al.^24^ illustrated that cancer-associated fibroblast-derived PAI-1 promotes the EndoMT in lymphatic ECs from cervical squamous cell carcinoma. Here, we identified that DHH directly interacts with PAI-1 and downregulate PAI-1 protein expression in a dose-dependent manner. This complex interaction between DHH and PAI-1 pinpoints DHH mediated negative regulation of PAI-1-induced pathophysiological effects, thereby offering protection against atherosclerotic cardiovascular diseases.

Notably, the hedgehog signaling has been extensively studied as a crucial regulator of cardiovascular development and maturation.^54–56^ Nonetheless, its role in atherosclerosis is controversial due to contradictory findings in the literature. For instance, research by Beckers et al.^57^ reported that inhibiting hedgehog signaling with an anti-Hh antibody aggravated atherosclerosis in *ApoE* KO mice due to the increased lipid uptake by macrophages. Conversely, a study by Zhang et al.^58^ has shown that blocking hedgehog signaling with the Hh pathway antagonist vismodegib ameliorated atherosclerosis and foam cell formation in *ApoE* KO mice by promoting autophagy in early atherosclerosis. These contradictory findings underline the complexity of hedgehog signaling and its context- and stage-dependent role in atherogenesis.

Our study adds another layer to this narrative, showing that *DHH* inhibits EndoMT possibly independent of canonical hedgehog signaling, as HAECs treatment with canonical hedgehog signaling inhibitor sonidegib didn’t alter *DHH* knockdown-induced or *DHH* overexpression-prevented EndoMT. A possible explanation for these findings is that *DHH* may mediate its effects through noncanonical pathways. Additionally, the discrepancies in the literature could stem from differences in experimental designs, including the stage of atherosclerosis being investigated, the cellular context (e.g., ECs, SMCs, or macrophages), or the specific pathways activated in response to hedgehog signaling.

We recognized that the present study has several limitations. First, the precise mechanism by which *Dhh* inhibits atherosclerosis development, whether via canonical or noncanonical hedgehog signaling *in vivo*, remains unclear. Therefore, it warrants further studies on whether EC-specific deletion of smoothened phenocopy *Dhh*^ecKO^ mice. Second, although *DHH* is highly expressed in ECs, we cannot exclude the contribution of VSMCs and macrophage-derived *DHH* to atherosclerosis under diseased conditions. Thus, studies using macrophage and VSMC-specific conditional knockout mice are crucial to deciphering the precise role of *DHH* in these vascular cells in the context of atherosclerosis. Last but not least, the potential of DHH as a diagnostic and prognostic marker for predicting future cardiovascular events remains to be explored in large-scale human population studies. Future endeavors will be dedicated to discovering pharmacological boosters of endothelial *DHH* expression in treating cardiovascular diseases.

Overall, our findings highlight *DHH* as a compelling candidate for potential therapeutic intervention in atherosclerosis due to its endothelial-specific expression and its ability to inhibit EndoMT via trapping PAI-1. As demonstrated by our current data, pharmacological restoration of DHH levels using recombinant DHH protein significantly mitigated atherosclerosis progression and promoted plaque stabilization in hyperlipidemic mice. These results suggest that DHH-based therapies may complement existing lipid-lowering strategies by enhancing endothelial resilience. Furthermore, reduced DHH expression in atherosclerotic plaques provides a rationale for considering DHH not only as a therapeutic target but also as a potential prognostic biomarker, reflecting the lesion progression or even vulnerability.

Nonetheless, potential risks associated with *DHH* overexpression must be carefully considered before clinical translation. As *DHH* is involved in angiogenic processes, ectopic or prolonged overexpression may inadvertently stimulate neovascularization or disrupt vascular integrity in off-target tissues. Additionally, because DHH inhibits PAI-1, it could interfere with the physiological role of PAI-1 in fibrinolysis, potentially increasing the risk of hemorrhagic complications or impaired wound healing. Although our study showed that *DHH* overexpression does not significantly alter canonical Hedgehog signaling components, long-term systemic elevation my still pose a risk of unintended pathway activation. Therefore, endothelium-targeted delivery strategies for DHH mimetics will be essential to maximize therapeutic benefit while minimizing off-target effects thereby preventing vascular homeostasis.

In conclusion, this study demonstrates that *DHH* is a novel mechanoresponsive and resilience factor in ECs that protects against atherosclerosis by inhibiting PAI-1-dependent EndoMT via competing with PAI-1 binding to LRP1 and ensuing pro-EndoMT signaling. The findings pave the way for additional studies to gain further insights into the therapeutic potential of modifying *DHH* signaling genetically or pharmacologically to treat atherosclerosis and other occlusive vascular disorders. Overall, our findings suggest that DHH downregulation is linked to the development and progression of atherosclerosis, indicating its potential as both a biomarker and a therapeutic target in cardiovascular disease.

## Acknowledgements

We are grateful to members of the Weng/Xu laboratory for assistance in tissue harvesting and insightful discussions.

## Authors contributions

S.X., and J.W conceptualized the project. I.I., and Z.W., performed the experiments, generated and analyzed the data. I.I. wrote the original manuscript. I.I. H.J., M.S., Z.Z., and F.Z. generated and analyzed the data. Y.F., L.W., Y.H., M.A.R., J.P., G.G.C., H.J., P.C.E., S.O., C.L.M., F.B.H.Z., S.W.L., L.M., Y.Z., Q.W., J.Z., B.Z., and T.J. made a substantial contribution to the analysis, generation of transgenic mice, and interpretation of study data and critiqued the manuscript for important intellectual content. All authors approved the manuscript.

## Supplemental Material

Expanded Materials and Methods

Figures. S1 to S21

Table S1

Key Resource Tables

Reference 59-76

Uncropped Gel Blots

Author Checklist

## Nonstandard Abbreviations and Acronyms

AA: Aortic arch
BCA: Brachiocephalic artery
CAD: Coronary artery disease
CHO: Cholesterol
DF: disturbed flow
EC: Endothelial cells
ecKO: Endothelial cell specific knockout
EndoMT: Endothelial to mesenchymal transition
HAEC: Human aortic endothelial cells
HUVEC: Human umbilical vein endothelial cells
LCA: Left carotid artery
PCL: Partial carotid ligation
TA: Thoracic aorta
TG: Triglycerides
UF: Unidirectional laminar flow

